## Supplemental Figures for "The sleep quality- and myopia-linked PDE11A-Y727C variant impacts neural physiology by reducing catalytic activity and altering subcellular compartmentalization of the enzyme"

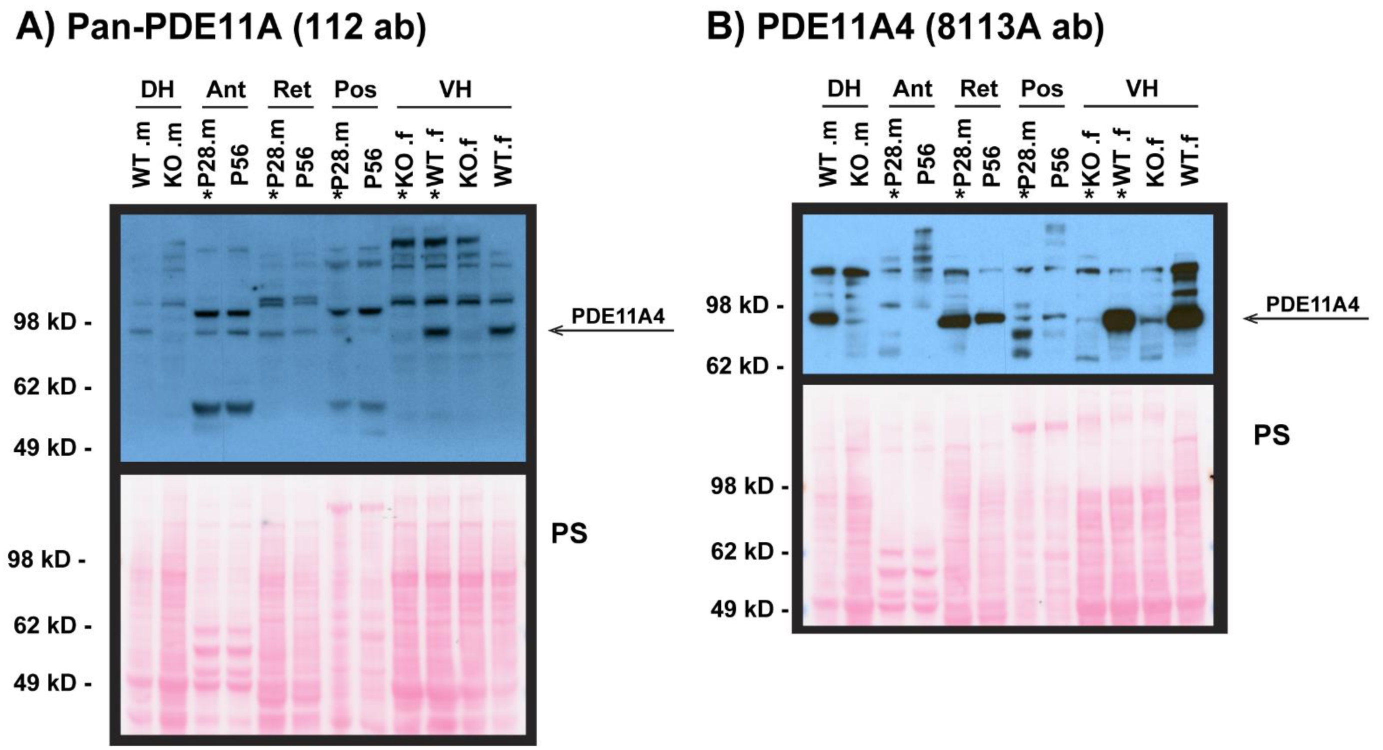


**Figure S1.** Pilot study Western blots confirm that acetone precipitation does not interfere with PDE11A4 signal detection. Dorsal (DH) and ventral hippocampal (VH) tissue is used as positive (WT) and negative (KO) controls. Samples that were treated with acetone are marked with (*). Ponceau stain (PS) images are provided as the loading control. PDE11A4 signal is detected in the anterior segment (Ant), retina (Ret), and posterior segment (Pos) tissues, using **A** the pan-PDE11A 112 ab and **B** the PDE11A4-specific 8113A ab. Brightness and contrast of blots and Ponceau stain (PS) images adjusted for graphical clarity. f – female; m – male.

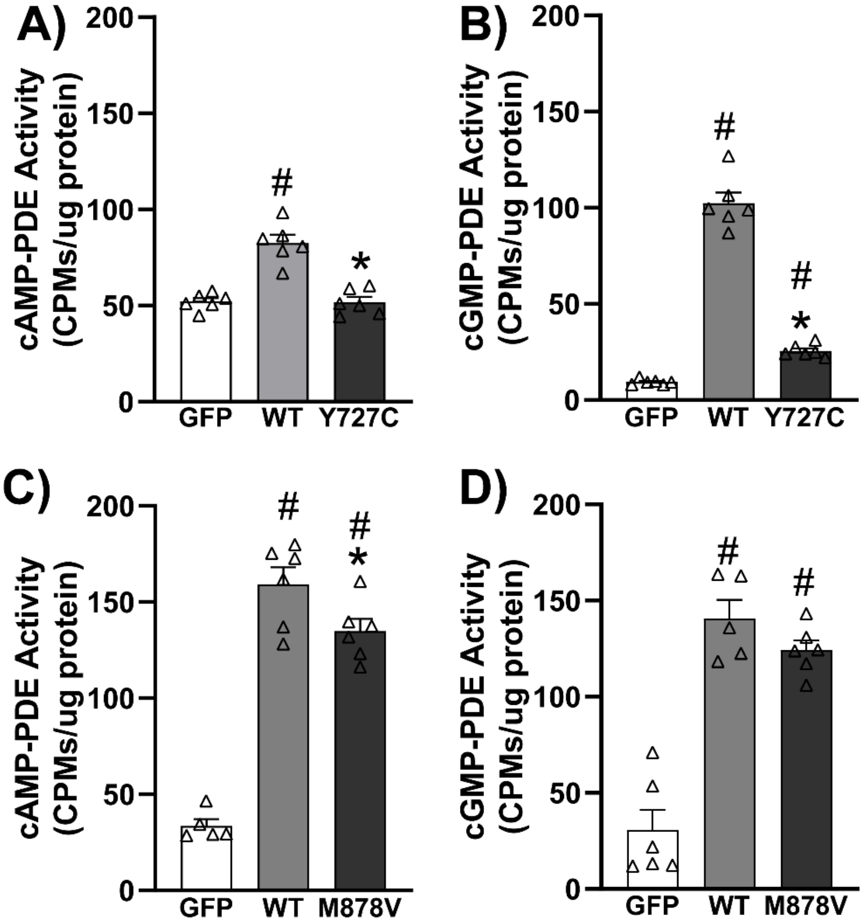


**Figure S2**. Replication of differential effects of Y727C vs M878V on cAMP- and cGMP-PDE11A4 activity (n=6 biological replicates/group). A) Y727C eliminates cAMP hydrolysis (F(2,15)=33.04, P< 0.0001; Post hoc vs 727C: WT P=0.0002, GFP P=0.9316). B) Y727C strongly impairs cGMP hydrolysis (failed normality H(2)=15.16, P=0.0005; Post hoc vs Y727C: WT P=.15, GFP P=.15). C) M878V impairs cAMP hydrolysis to a lesser degree than Y727C (F(2,14)=87.91, P<0.0001; Post hoc vs M878V: WT P=0.0225, GFP P=0.0002). D) M878V also impairs cGMP hydrolysis to a lesser degree than Y727C (F(2,14)=47.97, P<0.0001; post hoc vs M878V: WT P=0.21, GFP P=0.0002). Post hoc: *vs WT, P=0.0225-0.0002; #vs. GFP, P=0.0036-0.0002.

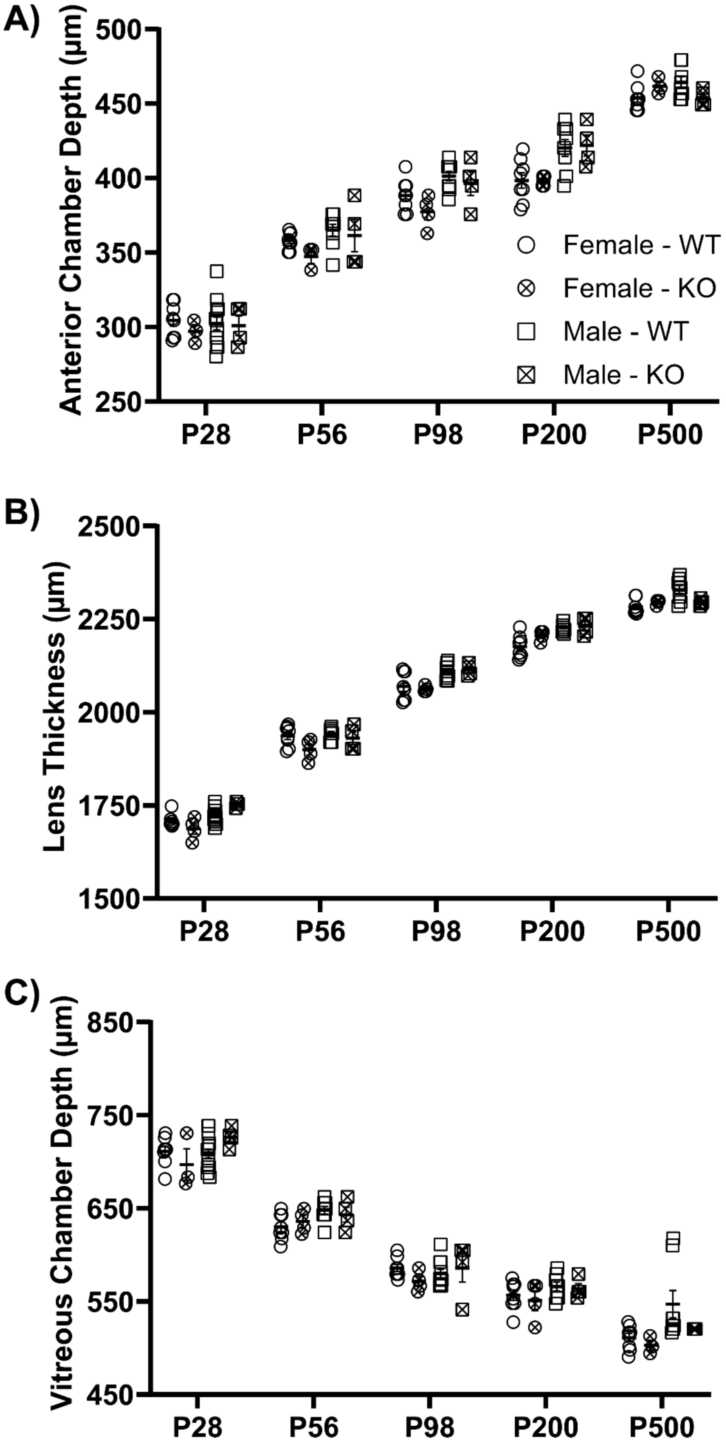


**Figure S3.** Ocular compartment sizes of Crispr B6/N *Pde11a* KO and WT male and female mice do not differ across the lifespan. A) Anterior chamber depth, B) lens thickness, and C) vitreous chamber depth changed with age but were not significantly different between *Pde11a* KO eyes (n=3-4 eyes from 2F + 3-4 eyes from 2M per age) and *Pde11a* WT eyes (n= 8 eyes from 4 F at each age + 8 eyes from 4M at P56, P200 and P500, 13-14 eyes from 7M at P28, and 9-10 eyes from 5M at P98). Data expressed as mean ± SEM with individual data points plotted over (circles, females; squares, male).

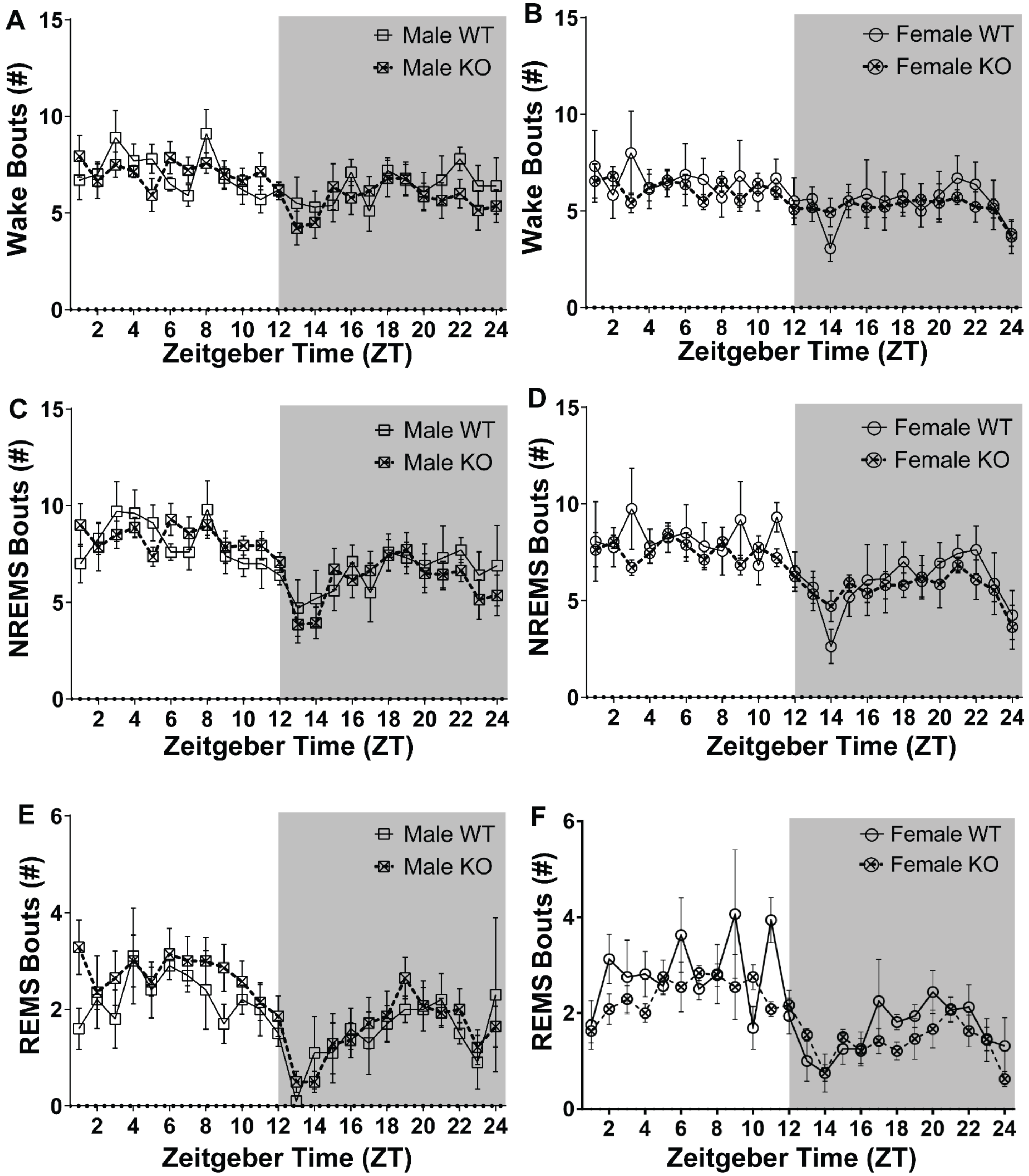


**Figure S4.** There are no significant differences between LacZ B6/J *Pde11a* KO and WT littermates on number of A-B) waking, C-D) NREMS, or E-F) REMS bouts.

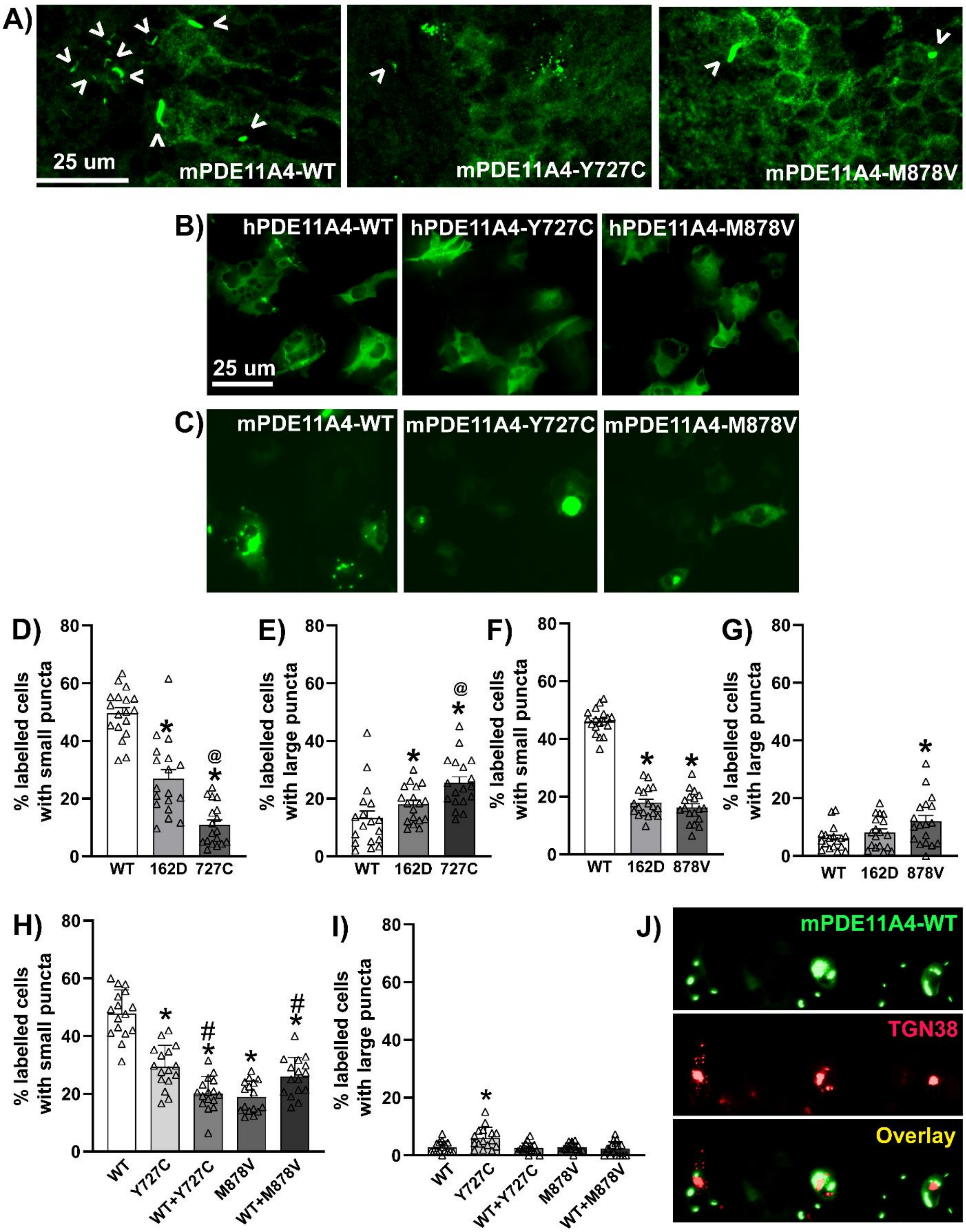


**Figure S5.** PDE11A4-Y727C and PDE11A4-M878V dispersal phenotypes observed in HT22 hippocampal cells were replicated in the male and female mouse hippocampus *in vivo* and in COS-1 monkey fibroblasts *in vitro*. A) Immunofluorescence for EmGFP-tagged PDE11A4 that was virally infused into the hippocampus of old male and female *Pde11a* KO mice (males shown) showing recombinant mPDE11A4-WT exhibits substantial punctate accumulation in linear filamentous structures termed "ghost axons" (indicated by arrowheads; [1]). In stark contrast, mPDE11A4-Y727C demonstrates a much more diffuse distribution, only rarely accumulating in linear ghost axons (and only to a minimal degree) but routinely accumulating in large clusters of circular

PDE11A4 puncta. mPDE11A4-M878V does not show these large accumulations of puncta, but does demonstrate a much more diffuse distribution pattern than mPDE11A4-WT, with accumulation in sparse ghost axons. B) Immunofluorescence of hPDE11A4 in COS1 cells and C) direct imaging of EmGFP-mPDE11A4 in COS1 cells shows accumulation of PDE11A4-WT, but dispersion of PDE11A-Y727C and PDE11A4-M878V. D) Quantification of mPDE11A4 shows that Y727C reduces the accumulation of small PDE11A4 puncta even more than does S162D (H(2)=37.78, P<0.0001; Post hoc vs Y727C: WT P<0.0001, S162D P=0.0142) while E) increasing the presence of large mPDE11A4 accumulations relative to WT (H(2)=16.47, P=0.0003; Post hoc vs WT: Y727C P=0.0002, S162D P=0.1958). F) mPDE11A4-M878V also reduces mPDE11A small puncta in COS-1 cells like mPDE11A4-S162D (F(2,51)=205.09, P<0.0001; Post hoc vs M878V: WT P=0.0001, S162D P=0.315). In contrast to HT22 Cells, however, G) M878V increases the presence of large mPDE11A4 accumulations in COS-1 cells (F(2,51)=4.10, P=0.0224; Post hoc vs WT: M878V P=0.0192, S162D P=0.3652). H) Also in contrast to HT22 cells, COS1 expression of WT and Y727C in equal proportions actually amplifies the dispersing effect of the variant on small PDE11A4 puncta relative to Y727C alone (F(4,75)=47.16, P<0.0001; Post hoc vs Y727C: WT P=0.0001, Y727C+WT P=0.0008), I) but completely blocks the effect of Y727C on increasing large PDE11A4 accumulations (H(4)=14.50, P=0.0059; Post hoc vs. Y727C: WT P=0.027, Y727C+WT P=0.0234). B-G, n=18 biological replicates/group over 3 experiments. J) Imaging of GFP-tagged mPDE11A4 and RFP-tagged TGN38 shows that small PDE11A4 puncta often appear to bud off the Golgi but never colocalize (see cells on left and right of image; as previously described [1]); however, large PDE11A4 accumulations can partially co-localize (see cell in center). *Post hoc*: *vs WT, P=0.049 to <0.0001; @vs S162D, P<0.05.


1. Pilarzyk, K., et al., *Conserved age-related increases in hippocampal PDE11A4 cause unexpected proteinopathies and cognitive decline of social associative memories.* Aging Cell, 2022. **21**(10): p. e13687.
